# A Fine-tuned ProtGPT2 (transformer model) for Predicting more Virulent SARS-CoV-2 variants and understanding its virulence by biophysical methods

**DOI:** 10.1101/2025.01.13.632691

**Authors:** Sam Paul Dhamodharan, Abraham Rebairo Jeyaraj, Rashmi Siddalingappa, Rajakannan Venkatachalam, Gugan Kothandan

**Affiliations:** CAS in Crystallography and Biophysics, University of Madras, Chennai, India; Department of Nuclear Physics, University of Madras, Chennai, India; Department of Computer Science, Christ University, Bangalore, India

## Abstract

The emergence of Variants of Concern (VOCs) of SARS-COV2 with increased virulence and transmissibility has been linked to multiple mutations in the RBD region, altering their antigenic properties. In this study, we used a specialized ProtGPT2 model trained on the SARS-COV2 spike protein to forecast probable mutations on the spike protein that have not yet emerged. Upon prediction, we systematically studied the stability of single-site and multisite mutations using unbiased molecular dynamics simulations. Binding free energies were used to study the physicochemical significance of the mutations and their affinity to human ACE2 receptor. Our results show that there are specific hot-spots that mutate in the spike protein that enhance binding affinity through electrostatic and improved non-bonded interactions and highlight the role of specific energetic contributions in viral adaptation and infectivity. Our analysis revealed that the reduction of a disulphide bridge within sites 480-488 lowered the binding free energy and increased the flexibility of the loop region, enhancing its interface interaction with ACE, leading to a more virulent variant than Omicron.

## Introduction

Nearly 33% of the deaths attributed annually to identified infectious diseases are due to viruses. Viral infections occur when the virus’s spike glycoprotein attaches to and enters a host cell receptor [1, 2]. Enveloped viruses, such as influenza hemagglutinin and the envelope glycoproteins of HIV-1 and Ebola, utilize comparable spike-like proteins for cell infection [3, 4]. Following the advent of SARS-CoV-2 in 2019, over 7 million people have succumbed to the virus. [5] For SARS-CoV-2, the spike protein targets the Angiotensin-Converting Enzyme 2 (ACE2) on cell surfaces. [6–9] Mutations within the Receptor Binding Domain (RBD) of the SARS-CoV-2 spike protein directly affect its binding to ACE2 and enable antibody resistance [10, 11]. The emergence of Variants of Concern (VOCs) with increased virulence and transmissibility has been linked to multiple mutations in the RBD region, altering their antigenic properties [12–16]. The Omicron strain features around 30 mutations in its spike protein that enhance its ability to bind to the ACE2 receptor [17–19]. XBB-descended variants in dominance by 2023 carry 45–48 mutations in the spike protein [20]. The World Health Organization (WHO) has categorized new and circulating SARS-CoV-2 variants into variants of concern (VOC), interest (VOI), and under monitoring (VUM) [21]. VOCs are notably aggressive, more transmissible, and resistant to vaccines, thus leading to severe infections. Many VOCs have emerged due to mutations in the spike protein of SARS-CoV-2 that increase binding affinity with human ACE2 receptors, leading to greater transmissibility [**?**]. WHO-designated VOCs include Alpha (B.1.1.7), Beta (B.1.351), Gamma (P.1), Delta (B.1.617.2), and Omicron (B.1.1.529), with Omicron being the most prevalent for nearly three years [22, 23]. Despite global vaccination efforts, the development of new variants has resulted in immune evasion [24]. The virus continually evolves, adapting to natural immune responses and vaccines (Han et al. 2023). SARS-CoV-2’s infection mechanism and mutation patterns provide critical insights into viral mutation processes overall. Specific mutations in the spike-proteins such as D614G [25], N501Y [26], E484K [27], T478K [28], and K417N/T [29] are known to increase infectivity, transmissibility, and immune evasion. These mutations demonstrate the complex physiochemical interactions influencing the spike-protein’s binding to the ACE2 receptor, both within and outside the RBD region. To this end, artificial intelligence (AI) based techniques has revolutionized many areas of computational Biology. [30–33] Machine learning (ML) models are quick and efficient in identifying patterns and relationships in the available vast amounts of biological data. Many of these techniques make use of deep learning, a branch of machine learning that uses neural networks to draw conclusions from vast volumes of data. In the case of SARS-CoV-2, genome data available, since 2019, is vast and deep learning models are being used efficiently to predict and understand the evolution of viruses. [34] ML models have been developed to identify persistent variants that remain over a period of time [35]. Domain-specific mutations are predicted by a ML model using entropic information of the spike protein. A risk assessment ML model is being helpful in forecasting new wave of infection in the following months [36]. The VPRE (viral protein evolution prediction) model forecasts viral evolution up to five months into the future [37]. Apart from Deep Learning models, Natural Language Processing (NLP) techniques are also increasingly being utilized for sequence-based prediction tasks in proteins. [38–40] In protein language models (PLM), individual amino acids are treated as tokens. Transformer-based models have advanced the PLMs by providing sophisticated approaches for sequence prediction. These models, such as BERT (Bidirectional Encoder Representations from Transformers) [41], utilize bidirectional masked language modeling to predict masked portions of the sequence. BERT’s Transformer architecture includes both LSTM and attention mechanisms [42], enabling it to handle complex sequence dependencies. MutFormer [43] an adaptation of BERT, incorporates an adaptive vocabulary for protein contexts to predict the impact of missense mutations on protein function. Other Transformer-based models like TEMPEL [44] and TEMPO [45] also contribute to predicting protein mutations. ProtBERT [46], a deep language model for protein sequences, captures local and global representations of proteins in a natural way. ProtTrans [47] helps understand protein properties. In this article we have used a Fine-tuned ProtGPT2 model to predict the mutations in the spike proteins of SARS-CoV-2, and thereby report more infectious variants. ProtGPT2 [48], is a transformer-based model, and we have fine-tuned the model on SARS-CoV-2 spike protein sequences. ProtGPT2 was pretrained on huge number of protein sequences in UniRef database and predicts ‘next-residue’, similar to ‘next-word’ prediction by Large Language Models (LLMs). We have used the finetuned (FT)-ProtGPT2 to generate possible mutations that may occur in future in the spike region of SARS-CoV-2. Unlike some of the models mentioned above, FT-ProtGPT2 does not predict the time frame of the mutation predicted. The predicted mutations of FT-ProtGPT2 are such that may have never occurred already or can occur any time in future. The stability of the mutants was confirmed through molecular dynamics simulations. Using techniques from structural biology, we have studied the physico-chemical significance of the mutations predicted, along with the combinations of predicted mutations as well. In the methodology section, we explain the Fine-tuned ProtGPT2 model and protocols followed in Molecular dynamics simulations. In the Results and discussion section, we systematically study single point, pairwise and combinations of several point mutations predicted by Fine-tuned ProtGPT2 and show the emergence of more virulent variants.

## Materials and methods

ProtGPT2 is a transformer-based decoder-only model pre-trained on 48 million protein sequences in UniRef50 database (version 2021 04). The model was trained in a self-supervised fashion, i.e., the raw sequence data were used during training without including the annotation of sequences. ProtGPT2, trained using a causal modeling objective, predicts the next token (in this case, oligomers) in the sequence. ProtGPT2 could be used in a zero-shot fashion or to generate sequences of a particular type after fine-tuning on a user-defined dataset. For fine-tuning ProtGPT2 and predicting mutations in the RBD region of the spike protein, we have used a procedure adapted by Dhodapker [49]. Figure 2 shows the protocol to finetune ProtGPT2 using SARS-CoV-2 spike protein. The raw data was the surface glycoprotein sequences downloaded from NCBI National Library of Medicine database. [50]. All human surface glycoproteins with sequence length 1273 from December 2019 until December 2023 (Raw data) were curated by eliminating redundant sequences, retaining only unique sequences (processed data). The processed data was then split into training and test set to validate the prediction efficiency of the fine-tuned model. We validated ProtGPT2 using different sets of ‘processed data’ of spike protein sequences. To verify the prediction efficiency of the fine-tuned model, we tested three cases – one VOC and two VOIs. The three cases were – Omicron (VOC), JN.1 (VOI), and XBB.1.5 (VOI). For example, the earliest sample date for Omicron is 29 September 2021. Therefore, to predict Omicron mutations (hereafter addressed as ‘Omicron case’) processed spike protein sequences before 29 September 2021 were only considered for train/test set (pre-Omicron) for fine-tuning ProtGPT2. From all the samples, 20,000 training samples and 10,000 test sequences were randomly selected for fine-tuning the model. The ProtGPT2 model was downloaded from the HuggingFace transformers model hub. While fine-tuning ProtGPT2, the hyperparameters that were adjusted accordingly. Since the spike protein residues between 406 and 506 form the receptor binding domain (RBD), we generated two sets of 1,00,000 sequences using two different prompts. Details on the sequences generated are explained in the Results and Discussion section. The synthetic amino acid sequences generated were aligned to the wild type (Wuhan SARS-Cov-2) spike sequence and mutations predicted by the model were identified using BioPython libraries [51]. All-atom molecular dynamics simulations were carried out for RBD mutants predicted by the fine-tuned model. We have used GROMACS v2022.3 for the study. The initial coordinates of the SARS-CoV-2 – ACE2 complexes were obtained from the protein data bank (PDB) with PDB IDs 6M0J and 7WHH corresponding to the wild type (WT) and Omicron complexes, respectively. Structures for spike mutants predicted by the model were prepared from WT using Chimera. The input complex structure in PDB format was converted into a GROMACS-compatible format using pdb2gmx.

**Fig 1.**
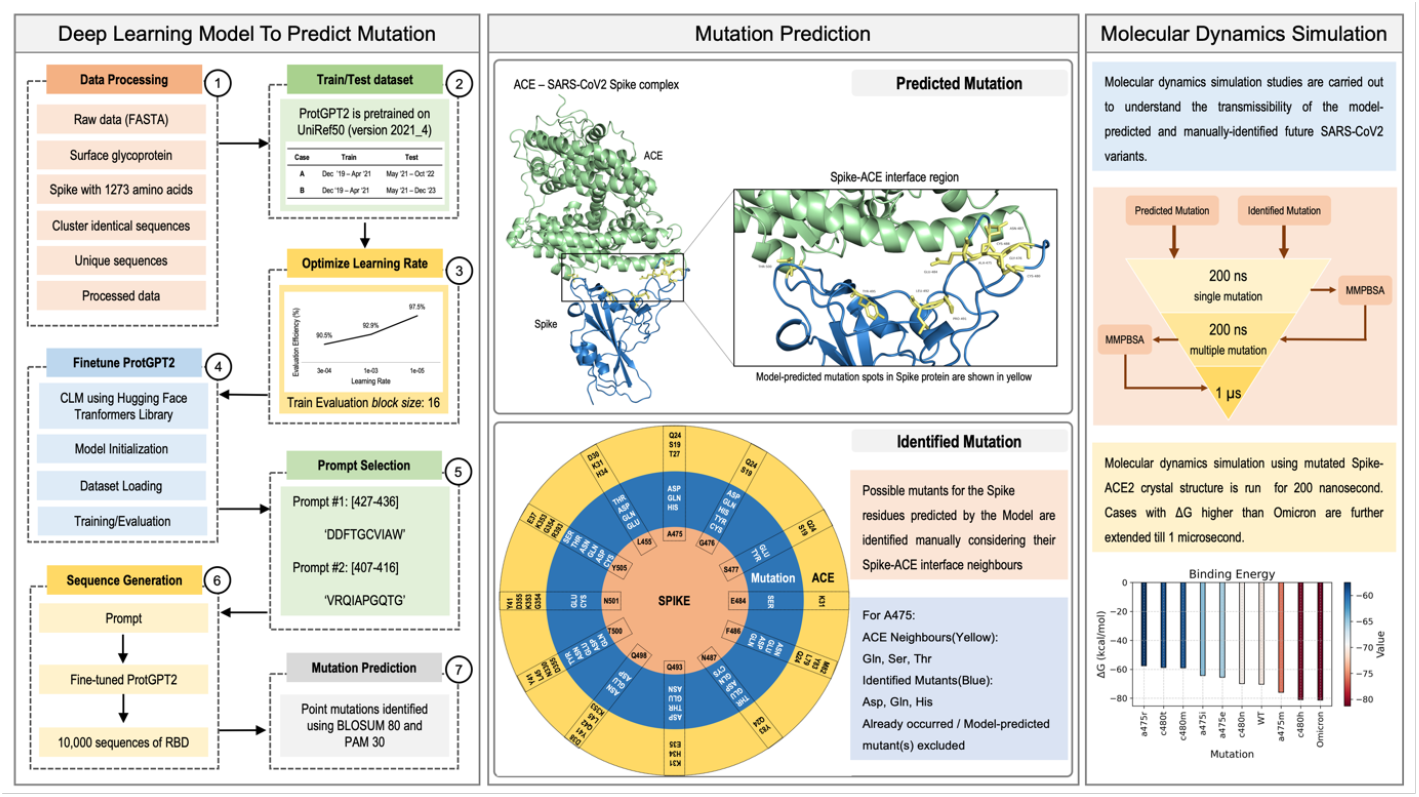
shows the scheme we have used to forecast the future variants of SARS-CoV-2 virus and to understand their binding with ACE2 receptor. The three main steps are: 1. Finetune ProtGPT2 deep learning model, 2. Predict mutation using Finetuned ProtGPT2, and 3. Molecular dynamics simulation and Binding free energy studies.

**Fig 2.**
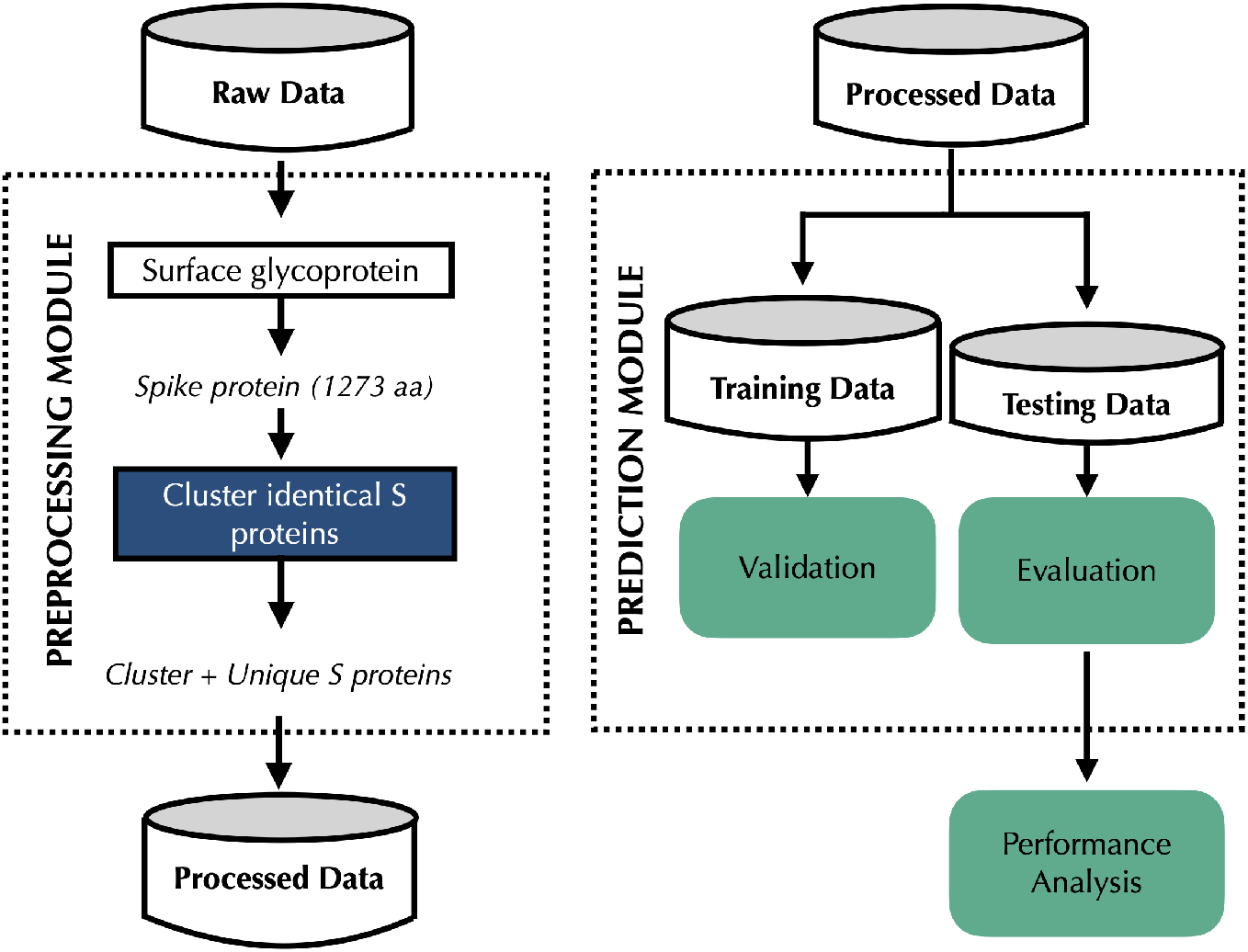
Protocol to fine-tune ProtGPT2 using SARS-CoV-2 spike protein

AMBER99SB-ILDN [52] force field was used for all the MD simulations reported in this work. The solute was centered in a cubical box with buffer between the boundary of the box and the solute set to 10.0 *Å*, and TIP3P water model was used. The solute-solvent interactions were neutralized through addition of sodium (Na^+^) counter-ions. The solvent molecules and counter-ions were initially relaxed by means of energy minimization. The system is gradually equilibrated to 300 K for 100 ps in NVT ensemble. V-rescale thermostat coupling method was used to adjust the temperature. Subsequently, the density was adjusted by means of 500 ps in NPT ensemble using Parrinello-Rahman barostat. The equilibrated structure was used for the production run. Variants with single point mutations, and combinations of several single point mutations were initially subjected to 200 ns unbiased simulations, and longer simulations upto 2 *µ*s for certain cases of interest. The bond lengths were constrained using LINCS algorithm [53]. The Verlet scheme was used as the cutoff-scheme for non-bonded interactions with a 1.0 nm set as cutoff for Coulomb and Van der Waals interactions. Long-range electrostatic interactions were treated using Particle-mesh Ewald (PME) method. All simulations were carried out on Linux operating system using NVIDIA A100 GPU card. Forty-three single, fourteen double, eight triple, seven tetra, four penta, three hexa, two hepta, and one octa mutation variants were studied in this work. The Chimera program was used to mutate residues in the spike protein and generate mutated variants. Few important cases were further extended to 1 *µ*s and also as triplicates. Using the obtained trajectories, Root-Mean-Square Deviation (RMSD) and Root-Mean-Square Fluctuation (RMSF) analysis were carried out using GROMACS tools to study structural stability of the mutated variants. Binding free energy with Poisson-Boltzmann (PB) solvation model and binding free energy decomposition were calculated using *gmx MMPBSA* [54]. The trajectory was divided into frames and binding free energy was calculated for each frame. For each snapshot i.e. each frame, the molecular mechanics energy (E_MM_) was calculated using AMBER FF14SB [55] force field, and the solvation energy (E_solv_) was calculated using Poisson-Boltzmann (PB) model. The binding free energy Δ*G*_*binding*_ was calculated,

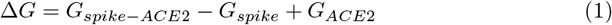

Where Δ*G*_*spike*−*ACE*2_ is the free energy of the spike-ACE2 complex, *G*_*spike*_ is the free energy of the spike protein, and *G*_*ACE*2_ is the free energy of the ACE2 protein. The per-wise scheme of *gmx MMPBSA* provides information about pairwise interactions between residues.

## Results and Discussion

The spike protein is highly susceptible to mutation due to its central role in viral entry into host cells and its importance in neutralizing antibodies. It is a glycoprotein comprising the S1 (receptor binding domain) and S2 (membrane fusion) domains. Our study focused on examining mutations in the S1 (RBD) domain, as it is in the interface of ACE2. The validation of Fine-tuned ProtGPT2 was carried out using various processed sets of spike protein sequences from the S1 domain. To evaluate the model’s prediction accuracy, tests were conducted using three cases: Omicron (VOC), JN.1 (VOI), and XBB.1.5 (VOI). The first instance of reporting Omicron variant was on 29 September 2021. Consequently, ProtGPT2 was fine-tuned using processed data comprising spike protein sequences collected before this date for Omicron. We have used 20,000 randomly chosen training samples along with 10,000 test sequences to refine the model. Two prompts were used to generate synthetic sequences in the receptor-binding motif (RBM) region: Prompt #1: DDFTGCVIAW and Prompt #2: VRQIAPGQTG. The amino acid sequence DDFTGCVIAW represents the last 10 residues immediately before the RBM of the spike glycoprotein, spanning from position 438-506. VRQIAPGQTG represents the 10 residues preceding the RBM, covering positions 406-495, including K417 of the RBD, which interacts with ACE. The validation studies was carried out with the Omicron case. For testing, both training and test sets contained sequences existing before the emergence of Omicron. A total of 100,000 sequences were generated per prompt. Fig. 3A presents the prevalence-weighted variant density estimates for the Omicron case. Previously identified mutations observed before the Omicron variant were included in the train and test sets. The ‘Predict’ section reveals novel mutations. Notably, the train and test distribution show relatively consistent mutation densities with minor peaks, whereas the ‘Predict’ section demonstrates sharper and more significant peaks, primarily between residues 470-510. High occurrences of novel mutations are found around residues 485-505, aligning with peaks observed in the ‘Predict’ section. Residues between positions 470-510 appear to be hotspots for mutations, especially novel ones, suggesting potential biological significance linked to the spike protein’s functional or structural roles. In a similar analysis, the Omicron case was evaluated using an alternative prompt, VRQIAPGQTG. The result depicted consistent densities with smaller peaks distributed among residues 405-495 (Figure shown in Supporting Information SF1). The Predict panel exhibited sharp, prominent peaks, notably around residues 475-495, with a marked increase in density toward the conclusion of this segment. Residues 490-495 showcased the highest prediction density. The residues within 470-495 indicated a significant increase in mutation counts, with a substantial amount categorized as ‘novel.’ Residues 485-495 exhibited the highest number of novel predictions, corresponding with the pronounced peaks in the Predict panel. This region, 470-495 showcased a high occurrence of anticipated mutations, suggesting its potential biological or evolutionary importance. Figures 3B and 3C illustrate the outcomes of JN.1 and XBB.1.5 test cases, prompted with prompt #1. Variants of concern identified include Alpha (September 2020), Beta (July 2020), Gamma (January 2021), Delta (October 2020), and Omicron (November 2021). ProtGPT2 was pretrained using the UniRef50 dataset (version 2021 04), utilizing spike protein sequences gathered before May 2021 for training. The FT-ProtGPT2 model’s efficacy in predicting new variants was assessed for the Variant of Concern (VOC) Omicron, and Variants of Interest (VOI): XBB.1.5 and JN.1, yielding 97%, 94%, and 94% evaluation accuracy on next-residue prediction using the test set of spike protein sequences for Omicron, XBB.1.5, and JN.1, respectively. Figure 3D presents the mutated amino acid (AA) positions in the receptor binding motif (RBM) of the spike protein, contrasted with the SARS-CoV-2 wildtype, for the three variants: Omicron, XBB.1.5, and JN.1. For XBB.1.5, which was first detected in September 2021, spike protein sequences collected post-May 2021 and pre-September 2021 were selected as the testing set. Analogous selection criteria were applied for Omicron and JN.1. Figure 3D illustrates the potential RBM mutations predicted by FT-ProtGPT2 for these three variants. Care was taken to ensure FT-ProtGPT2’s predicted mutations did not appear in either the training or testing datasets. Predicted variants are tabulated in Table 1, demonstrating that all mutations are unique. *Stability of mutations predicted by FT-ProtGPT2* MD simulations were done for both the wild type (WT) and Omicron variant up to 200 ns. Of the mutations predicted by FT-ProtGPT2, the mutations that have not occurred so far have been considered as future variants and were taken up for our study. In this study, forty-three single, fourteen double, eight triple, seven tetra, four penta, three hexa, two hepta, and one octa mutation combinations were performed. Molecular Dynamics (MD) simulations were executed for all single-point mutation variants over 200 nanoseconds. Figure 4A displays a heatmap of the predicted binding free energies for these mutations, as derived from FT-ProtGPT2 predictions. The binding free energies were calculated using the MMPBSA model. Among the mutations depicted in Fig. 4, eight (A475M, C480H, E484I, G476T, L492I, P491R, T500R, and Y495V) exhibit spike-ACE2 binding free energies that are lower than that of the Omicron variant (Δ*G* = −81.28 kcal/mol). Figure 4B presents the RMSD trajectories for these eight mutations across the 200 ns MD simulation. Analysis of the RMSD shows that C480H retains a considerable level of stability throughout the simulation, with RMSD values fluctuating between 2.0 −3.0*Å*, remaining closely aligned with WT. This indicates that the mutation does not significantly disrupt the protein’s stability and is unlikely to affect its overall function. Additionally, the stable RMSD of C480H correlates with a positive spike-ACE2 binding affinity, with a binding free energy of Δ*G* = −81.00 kcal/mol. Likewise, the RMSD for E484I also remains within 2.0-3.0 *Å* during the entire 200 ns simulation. Although there is minor structural rearrangement with the E484I mutation, as indicated by RMSD fluctuations between 50-150 ns, the structure stabilizes as the simulation progresses. The E484I mutation also demonstrated strong binding energy (Δ*G* = −83.77 kcal/mol), surpassing that of the Omicron variant. Another significant case is T500R, which presents a binding free energy of Δ*G* = −87.60 kcal/mol, lower than that of Omicron. The RMSD of these single-point mutations provides critical insight into how the residue at the mutation site impacts the overall flexibility of the spike protein. Figure 5A illustrates the root mean square fluctuations (RMSF) for the predicted point mutations, alongside WT and Omicron. RMSF represents residue flexibility during MD simulations, with peaks indicating highly flexible loop or surface-exposed regions. Residues 350-375 and 500-525 are identified as high-flexibility areas, while residues 425-450 are notably stable. C480H shows consistently lower RMSF in the rigid 425-450 region, reflecting its increased rigidity. Conversely, T500R demonstrates reduced RMSF in the flexible 500-525 region, which could enhance its stability.. We conducted a study focusing on the binding free energy contribution per residue for variants with an RMSD between 2.0 − 3.0*Å*. This involves the T500R, E484I, and C480H variants. The energy components per residue for C480H are depicted in Figure 5B, while those for E484I and T500R are displayed in Figures 5C and 5D, respectively. Figure 5B reveals that the Glutamic acid at position 484 significantly increases the binding free energy, attributed to its electrostatic repulsion with adjacent residues. The mutation from Cysteine to Histidine at position 480 disrupts a disulfide bond between positions 480 and 488, enhancing the flexibility of the loop area near the interface, though Glutamic acid’s energetic dominance persists throughout the simulation. For the E484I variant analysis, the Glutamic acid is replaced by Isoleucine, decreasing the region’s polarity and stabilizing the binding energy. Unlike the C480H case, Isoleucine does not increase binding free energy from position 484. In the T500R scenario, despite the mutation at position 500, Glutamic acid at 484 still largely influences the binding energy. Detailed structure examination indicated that binding energy is greatly affected by mutations between positions 480 and 500, in an unstructured loop near the ACE2 interface. Notably, previously studied variants E484K [27] and T478K [28] exhibited increased virulence, as both mutated Glutamic acid and polar uncharged side chains to lysine, which is positively charged. Their stability is attributed to complex many-body physico-chemical interactions. On taking the C480H case, the mutation involves reducing a disulphide bridge. The relatively stable RMSD of C480H throughout the 200 ns simulation correlates with the favourable spike-ACE2 binding affinity (Δ*G* = −81.00 kcal/mol). The di-sulphide bridge between the cysteine residues is present in the wild-type variant. The histidine residue at the 480^th^ site can possibly act as a redox center as it is close to the interface region. The histidine (480) – cysteine (488) interactions are primarily electrostatic in nature due to their partial charges in the sulphur of cysteine and nitrogen of the imidazole ring in histidine. On further investigation we found that there are multiple non-local interactions within the loop region, affecting the binding affinity. The point mutation at 484^th^ site from Glutamic acid to Isoleucine is also a significant prediction. A pairwise combination of mutations C480H and E484I is also done in order to check the stability of the binding, and shows binding energy in the range of wild-type variant (Δ*G* = −70.00 kcal/mol). The negatively charged Glutamic acid residue at 484^th^ site is weak to contribute to destabilizing histidine – cysteine interactions in 480 – 488 sites, whereas, when the disulphide bridge is reduced, the interaction becomes strong.

**Table 1.**
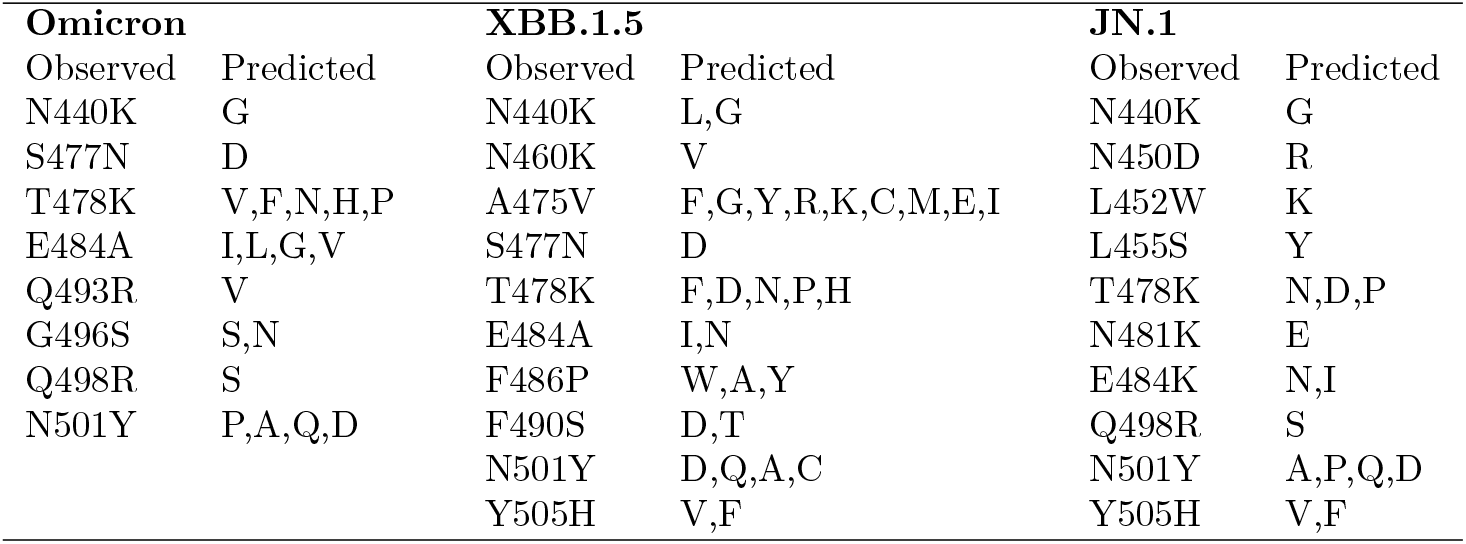
The actual mutations ‘observed’ in the variants Omicron, XBB.1.5 and JN.1 and the mutations ‘predicted’ by FT-ProtGPT2 are listed. G496S in Omicron was exactly predicted by the model.

**Fig 3.**
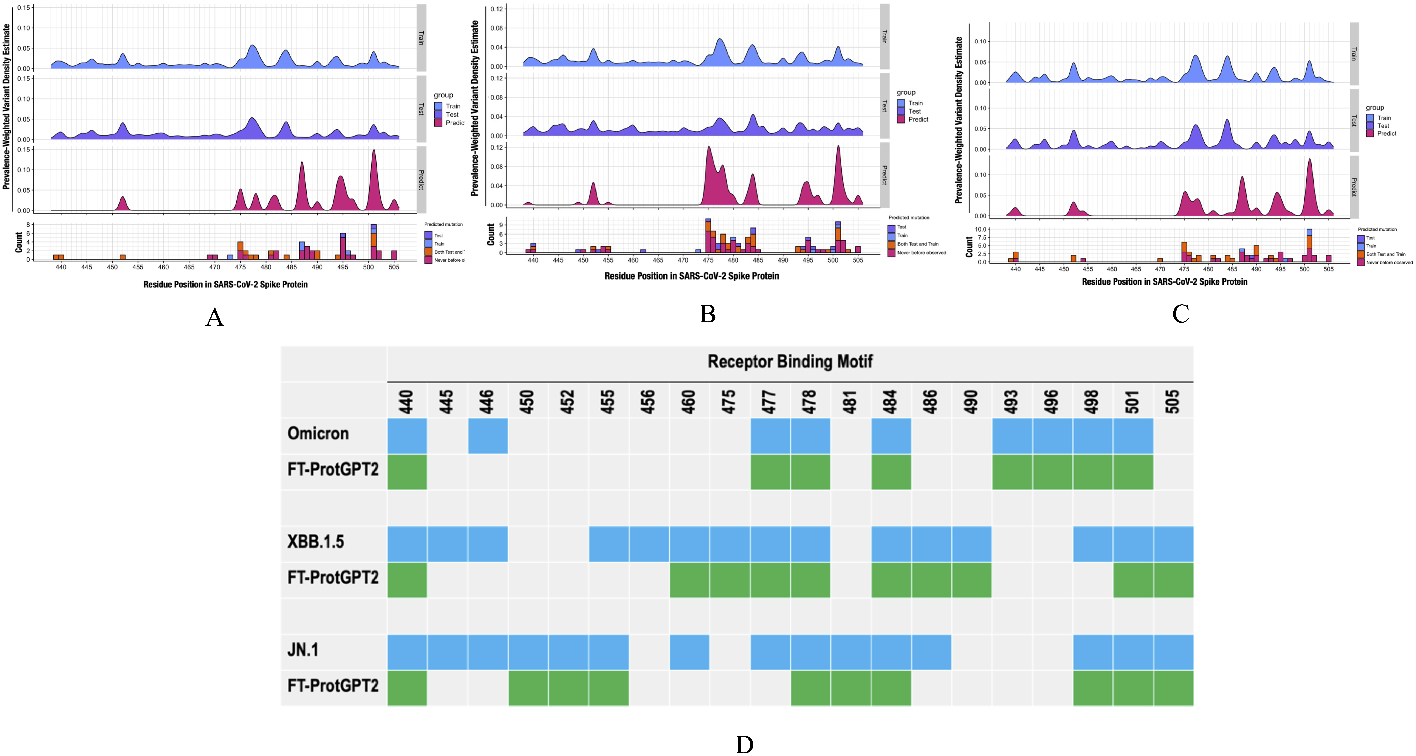
Prevalence Weighted density estimate of Omicron [426-515] (A), JN.1 [426-515] (B), XBB 1.5 [426-515] (C) variants and mutated amino acid (AA) positions of the receptor binding motif (RBM) of the spike protein (D) from those of SARS-CoV-2 wildtype for the three variants (Omicron, XBB.1.5, and JN.1) shown in blue and the probable mutations predicted by (FT-ProtGPT2) model in the RBM shown in green

**Fig 4.**
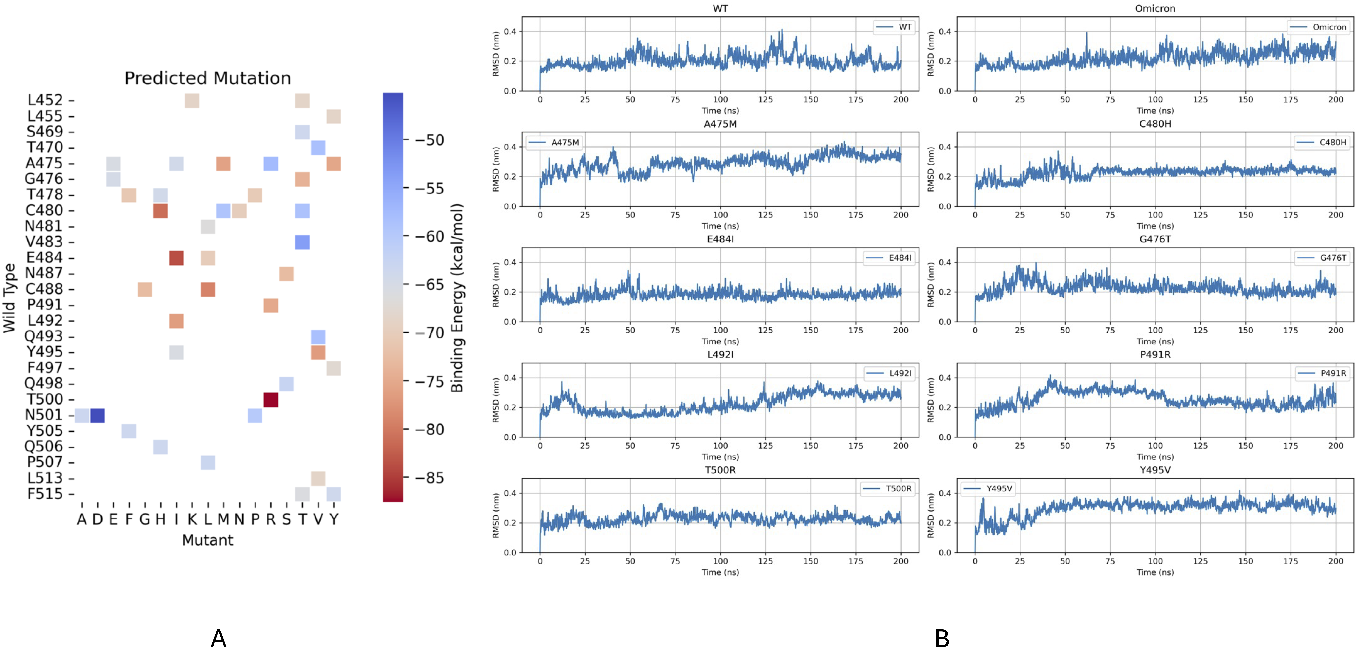
(A) Point mutation cases attempted through MD simulation for 200 ns. Wild type (WT) residues on the y-axis represent the position of the residues in the WT spike-ACE2 complex. The amino acids listed along with x-axis represent the mutations predicted by FT-ProtGPT2. Red shades indicate more favourable binding free energies. (B) RMSD trajectories of single-mutation cases for those with binding energy better than Omicron

**Fig 5.**
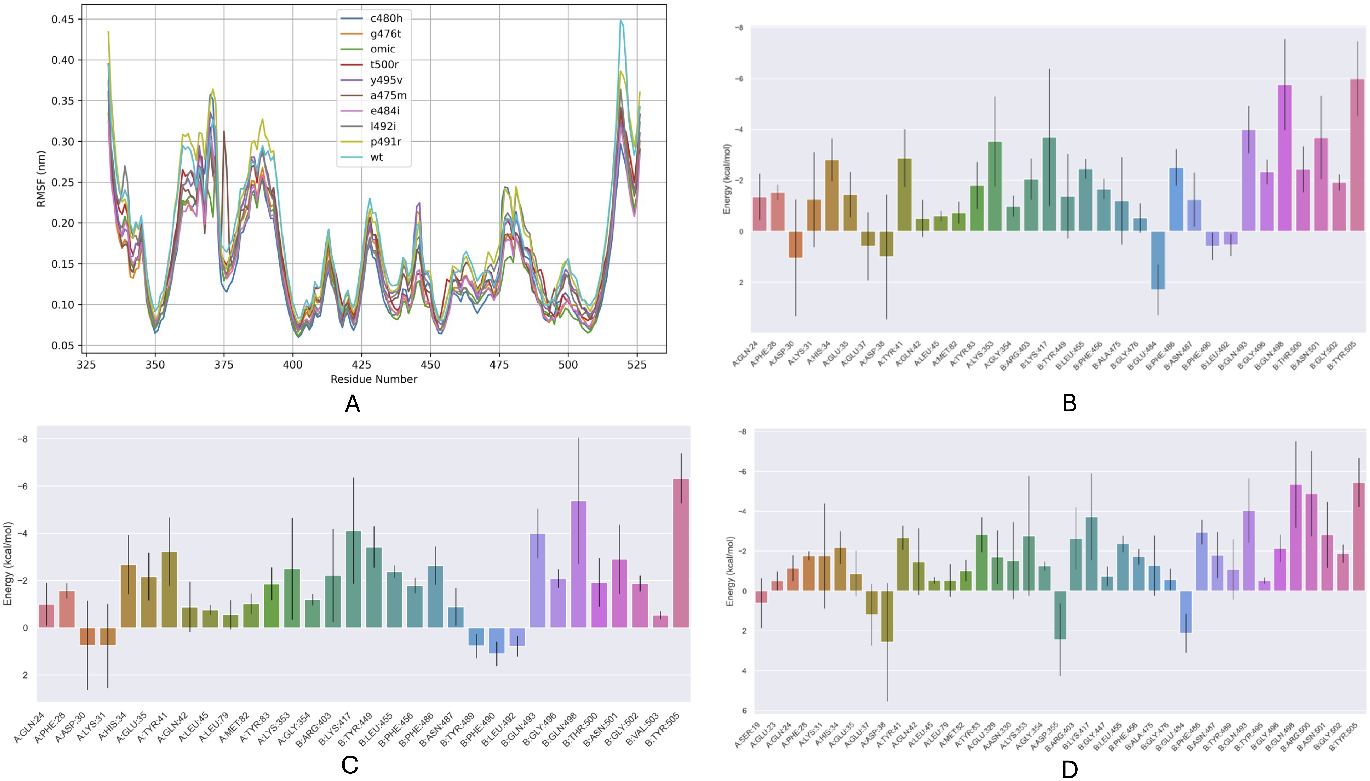
(A) RMSF of the Receptor binding motif for all the single point mutated variants. (B) Energetic components per-residue of C480H mutant (C) Energetic components per-residue of E484I mutant (D) Energetic components per residue of T500R variant

Mutation to a non-polar Isoleucine can in part reduce the electrostatic interaction in the region. As observed earlier, the overall stability of the spike-protein is unaffected from the RMSD trajectories (RMSD of E484I remains in the range of 2.0-3.0 *Å* throughout the 200 ns). E484I mutation showed strong binding (Δ*G* = −83.77 kcal/mol). Whereas in the T500R, the 500^th^ residue is mutated to Arginine from Threonine. This shows that a positively charged polar group, Arginine can form salt-bridge with negatively charged Glutamic acid of ACE. The formation of salt bridge post mutation occurs at the cost of reducing hydrogen bond interactions due to the removal of OH^-^ group from Threonine. We encounter similar patterns of reducing hydrogen bond interactions and neutralizing electrostatic interactions between heavy atoms in several point mutations predicted in this work. The shift from hydrogen bonds to salt-bridges enhances the binding specificity due to stronger interactions of salt-bridges relative to hydrogen bonds. The binding energy of each point mutations and their corresponding variations in the hydrogen bonds and non-bonded interactions in comparison to wild-type and Omicron variants is shown in Table 2. The NB/HB ratio shows the competing nature of hydrogen bond and non-bonded interactions. There is no direct correlation between the binding free energies and the NB/HB ratio, but it provides a good metric to identify stable interactions in certain variants. Variants with higher NB/HB ratios C480H with 4.70 and Y495V with 4.00 rely more heavily on non-bonded interactions compared to hydrogen bonds. This suggests that non-bonded interactions, specifically electrostatic interactions play a larger role in stabilizing these complexes. Variants with lower NB/HB ratios, A475M with 2.60 and G476T with 2.75, have a more balanced or hydrogen-bond-dominated interaction. This indicates that hydrogen bonding is critical in these mutants for ACE2 binding stabilization. WT has a moderate NB/HB ratio (3.14), indicating a fairly balanced contribution of non-bonded and hydrogen bond interactions. Omicron shows a low number of hydrogen bonds (9) and non-bonded interactions (28), but its NB/HB ratio (3.11) is close to WT. Despite fewer interactions overall, the non-bonded-to-hydrogen bond balance is preserved. G476T has 16 hydrogen bond (HB) and 44 non-bonded (NB) interactions, but its Δ*G* is not energetically significant. Whereas, Omicron with just 9 hydrogen bonds has relatively stable Δ*G* indicating few but more optimized bonds. C480H forms the most NB interactions (NB=47) leading to the stability of its binding. Omicron has the least NB interactions, yet achieves significant stability. This suggests that HB interactions alone don’t determine binding stability in this case, and that both hydrogen bonds and non-bonded interactions collectively contribute to stabilizing the protein-protein interface. Upon examining the hydrogen bond patterns throughout the entire trajectory for all point mutations, it becomes clear that electrostatic interactions (non-bonded) surpass hydrogen bonding in significance. This is evidenced in Figures 5B, 5C, and 5D, which present bar graphs of energy contribution per residue. From these figures, it’s apparent that Aspartic acid and Glutamic acid, present in T500R, C480H, and E484I, enhance their contribution to binding energy. These amino acids feature carboxylate groups and cause conformational shifts in the protein through long-range electrostatic interactions with positively charged groups. This overall dynamic interaction affects the protein’s binding stability. Further studies introduced combinations of several single-point mutations, termed multiple mutations. An MD simulation was conducted on variants with these multiple mutations, incorporating combinations of previously investigated single mutants that exhibited lower binding free energies than the Omicron variant.

**Table 2.** Number of hydrogen bond (HB) and nonbonded interactions (NB) between spike and ACE2 for the point mutation variants, sorted by increasing NB/HB ratio.

| | HB | NB | $\Delta G$ (kcal/mol) | NB/HB |
| --- | --- | --- | --- | --- |
| A475M | 15 | 39 | -75.94 | 2.60 |
| G476T | 16 | 44 | -74.01 | 2.75 |
| P491R | 15 | 46 | -75.14 | 3.07 |
| WT | 14 | 44 | -70.34 | 3.14 |
| Omicron | 9 | 28 | -81.28 | 3.11 |
| T500R | 11 | 36 | -87.60 | 3.27 |
| L492I | 11 | 39 | -76.45 | 3.55 |
| E484I | 10 | 39 | -83.77 | 3.90 |
| Y495V | 10 | 40 | -76.59 | 4.00 |
| C480H | 10 | 47 | -81.00 | 4.70 |

Figure 6 compares the binding free energies of WT, Omicron, and variants with multiple mutations, while Table S1 details the electrostatic and non-bonded contributions to the binding free energy for these variants. Notably, seven multiple mutation cases displayed better binding energy than Omicron, with 16 cases falling between the wild type and Omicron. For these seven cases, MD simulations were initially run for 200 ns. Several variants demonstrated lower binding free energies than the Omicron variant. Figure 7 analyzes the energetic components per residue of these variants compared to WT, Omicron, and Delta variants. According to Figure 6, variants with mutations at A475M-C480H-E484I-G476T-L492I-T500R and A475M-C480H-E484I-G476T-L492I-P491R-T500R-Y495V resulted in the lowest binding free energy, approximately 10 kcal/mol less than the Omicron variant. Evaluating the pairwise combinations of point mutations, the combinations N487S-C488I and A475M-C480H demonstrate the lowest binding energy, and in both cases, the disulfide bond is reduced due to mutation at the 480^*th*^ or 488^*th*^ sites. While these binding free energy shows the importance of loop region, and the role of di-sulphide reductions, several three-point mutations also show lower binding free energy by the mutation at 484^*th*^ site (A475M-E484I-N487S), which is also in the loop region as well. When we compared the energetic components per-residue of WT, delta, and Omicron with multiple mutations, we find that as long as site 484 mutates, the binding free energy lowers. This could be seen even in the WT, delta and Omicron variants in Figure 7A, 7B and 7C. The marginal reduction in the binding free energy in Figure 7E relative to 7A - 7D indicates the effect of electrostatic interferences of the Glutamic acid increasing the binding free energy. The mutations at 475, 480, 484, and 500th sites becomes very crucial in this study. *Importance of Disulphide reduction* From Figures 7G, 7H, and 7I, the energy components per residue affecting the binding free energy of various mutations are presented. In Figure 7G, the binding free energy is Δ*G* = −93.51 kcal/mol, while Figure 7H shows Δ*G* = −92.81 kcal/mol. These mutations were identified as the most advantageous according to FT-ProtGPT2s predictions. Notably, when the C480H mutation is excluded, as shown in Figure 7I, the binding free energy rises to Δ*G* = −83.18 kcal/mol, even though other point mutations remain unchanged. Further research indicates that breaking the disulfide bond enhances conformational flexibility in the loop region. This flexibility functions as a latch-release mechanism, encouraging more favorable interactions with the ACE2 receptor. We explored as to what makes the C480H mutation significant. Histidine differs from cysteine in side chain structure, which can modify the electrostatic environment at the binding site. The C480H mutation is crucial largely because it influences the protein’s redox state and electrostatic characteristics. Redox states are vital for protein structure and function, and altering redox activities can substantially change how proteins behave and interact. When cysteine is replaced by histidine at position 480 (C480H), the disulfide bonds cannot form, leading to more flexible protein structures. Moreover, losing a disulfide bond might impact the protein’s overall redox environment. Disulfide bonds are part of electron transfer processes, and their disruption might alter oxidative stress or redox signaling. Introducing histidine, with its imidazole side chain, can change local pH levels and result in new hydrogen bonds, thereby modifying the electrostatic potential at the binding site. Overall, the cysteine-to-histidine mutation introduces structural, redox, and electrostatic changes, profoundly affecting protein behavior, interactions, and functionality. This alteration likely improves the binding site interactions, decreasing binding free energy, and enhancing protein-receptor binding affinity. The variants with the lowest Δ*G* are a475m c480h e484i g476t l492i p491r t500r: Δ*G* = −93.51 kcal/mol and a475m c480h e484i g476t l492i p491r t500r y495v: Δ*G* = −92.81 kcal/mol. This implies that these multi-mutated variants achieve the strongest binding to ACE2 among those listed in Supporting Information Table S1. Van der Waals interactions are significant across all variants and increase with the number of mutations. Comparing the Wild Type (*V dW* = −87.49 kcal/mol) to a475m c480h e484i g476t l492i p491r t500r: (*V dW* = −102.32 kcal/mol), we observe enhanced nonbonded van der Waals interactions at the Spike-ACE2 interface. Intriguingly, the Electrostatic interactions (EEL) energy is noticeably lower in mutated variants compared to Wild Type (EEL = -709.22 kcal/mol) and Omicron (EEL = -1420.90 kcal/mol). The multi-mutated variant a475m c480h e484i g476t l492i p491r t500r (EEL= -1418.90 kcal/mol) displays more negative Electrostatic interactions, indicating that these interactions are significantly optimized in multi-mutated variants, contributing to binding stability. Electrostatic energy (EEL) is the chief contributor to improved binding energy in mutated variants, while VDW contributions acts supportively. Within the range of residues 406-506, there are typically two disulfide bonds that contribute to the stability of the protein’s structure by linking different parts of the polypeptide chain. Breaking only one of these bonds, such as with the C480H mutation, introduces a level of flexibility that may enhance specific protein functions while maintaining overall structural integrity. This selective breaking allows the protein to be more adaptable at certain regions without completely destabilizing its overall conformation. Such flexibility can be important for interactions at the binding site, making it easier for the protein to adjust its shape to fit more snugly with the ACE2 receptor, thereby enhancing binding affinity.

**Fig 6.**
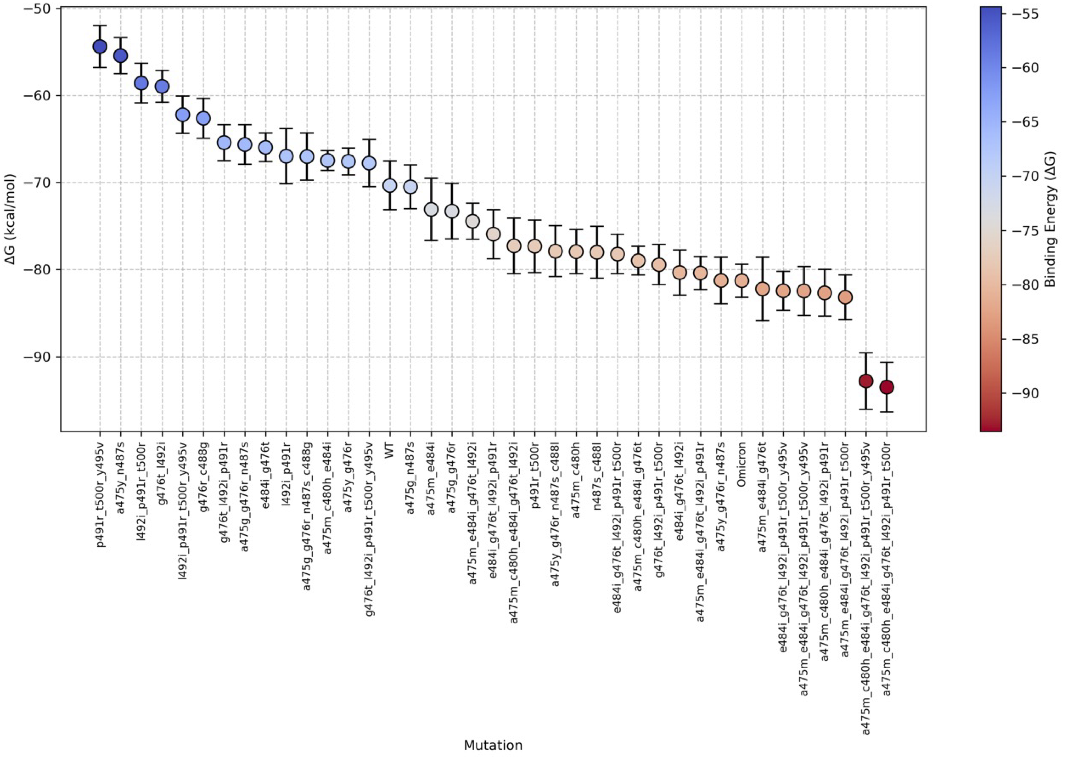
Binding free energy of WT, Omicron and FT-ProtGPT2 predicted variants with multiple mutations

**Fig 7.**
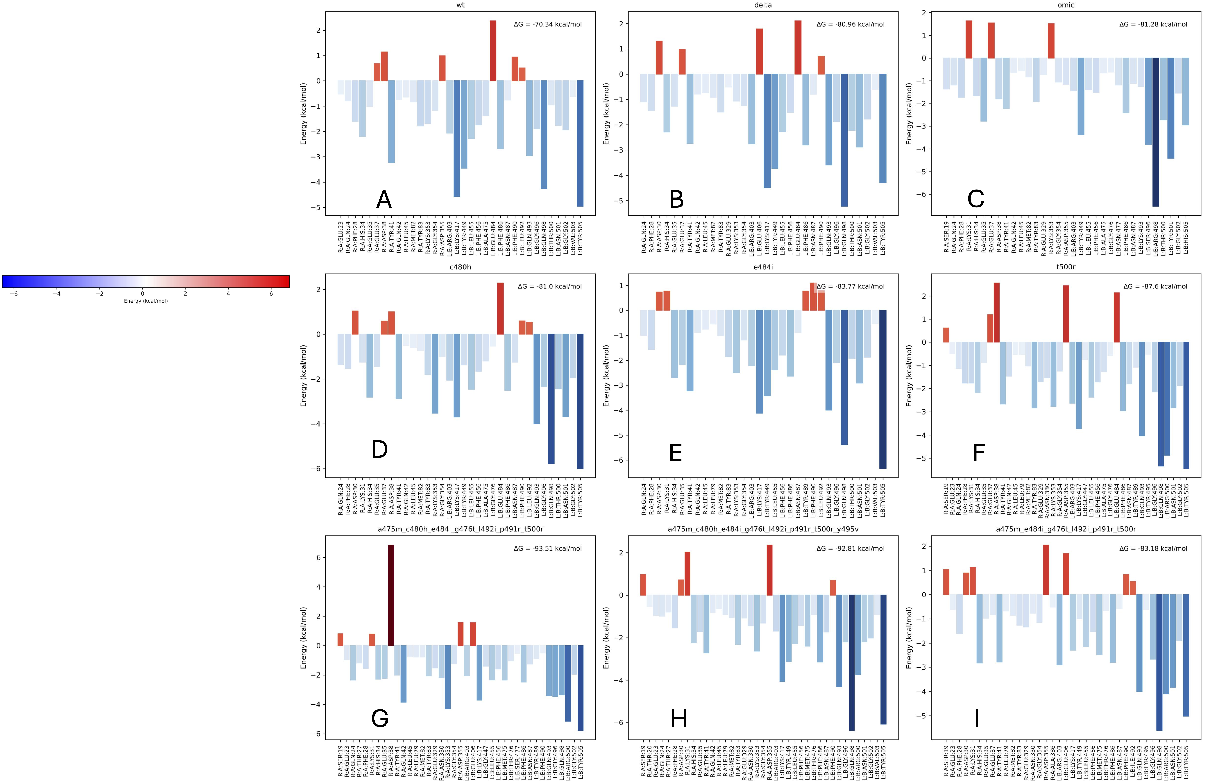
Energetic components per residue contribution to binding free energy of (A) WT (B) Delta (C) Omicron, variants with single point mutations (D) C480H, (E) E484I, (F) T500R, (G) A475M-C480H-E484I-G476T-L492I-P491R-T500R, (H) A475M-C480H-E484I-G476T-L492I-P491R-T500R-Y495V, (I) A475M-E484I-G476T-L492I-P491R

**Fig 8.**
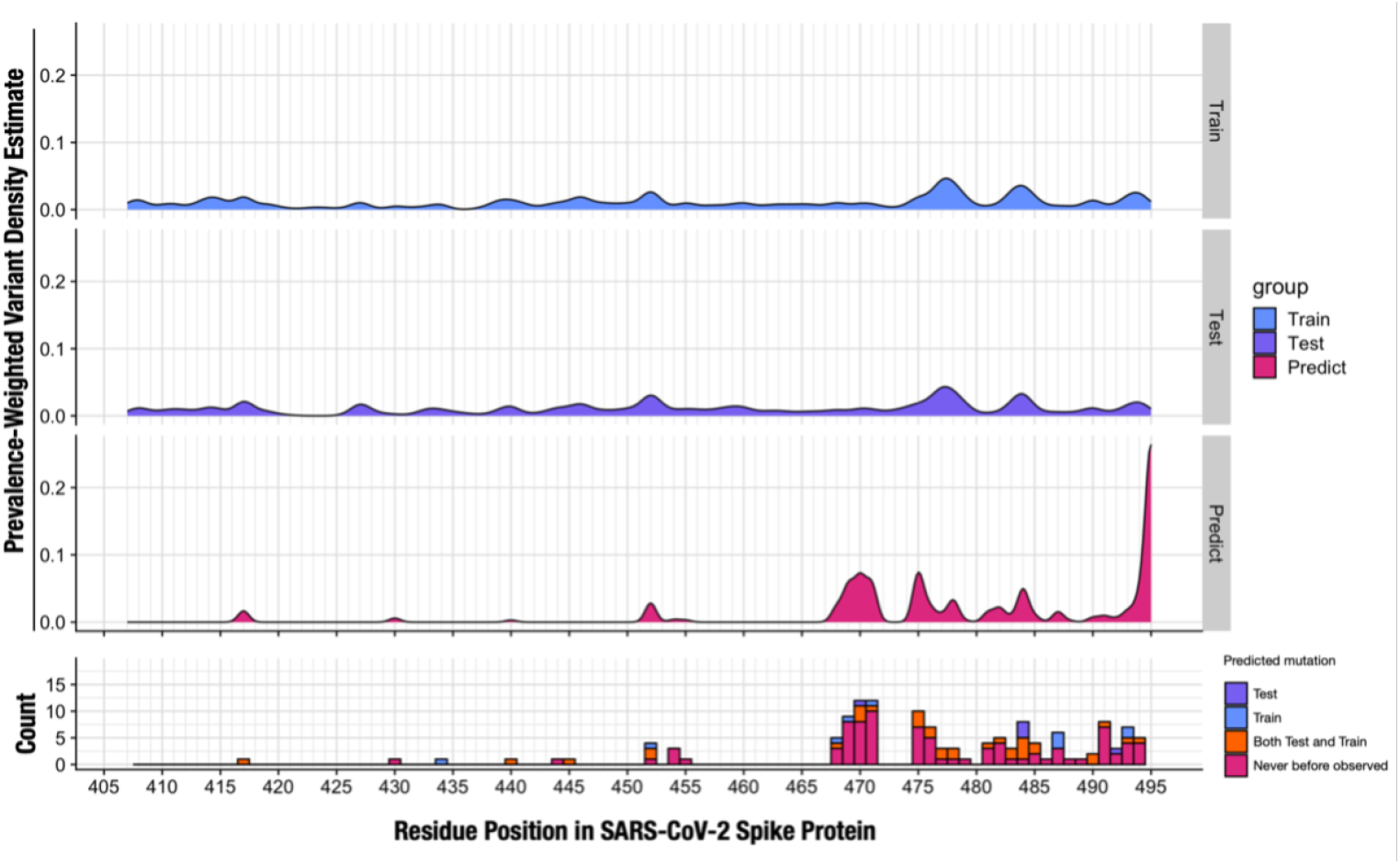
Prevalence Weighted density estimate of Omicron [405-495]

## Conclusion

In this study, a specialized ProtGPT2 model trained on spike-protein of SARS-CoV-2, Fine-tuned ProtGPT2 was utilized to predict new spike protein mutations in SARS-CoV-2 that have not yet emerged. This was done to gain further insight into the development of more virulent variants and to examine the physico-chemical effects of these mutations in the virus’s RBD. The training datasets incorporated the mutations reported in Alpha, Beta, Delta, and Omicron variants up to now. Mutational predictions were conducted using two different prompts, and their stability was tested through unbiased MD simulations. Additionally, the spike protein’s binding free energy with ACE2 receptors was explored at the residue level. By employing FT-ProtGPT2 to analyze mutations from single-point to pairwise and eventually multi-site, our findings highlighted a critical mutation-prone region in the spike protein (470-500). Further examination of the binding free energy per residue identified the loop region (478-488) as a hotspot for crucial mutations. Specifically, our analysis revealed that the reduction of a disulphide bridge within sites 480-488 lowers the binding free energy, resulting in variants more virulent than Omicron. Moreover, the combined mutations predicted by FT-ProtGPT2 further emphasize the role of disulphide bond reduction in decreasing binding free energy. These findings underscore the importance of the loop region (478-488) near the interface as a mutation hotspot and suggest that breaking a disulphide bond (Cysteine-Histidine) between sites 480-488 might contribute to the emergence of more virulent future variants.

## Acknowledgments

The Corresponding Author Gugan Kothandan would like to thank the Indian Council of Medical Research (ICMR) for funding this project (ID: 2021-15766).

## Supporting information

**Table S1.**

| Variant | VDW | EEL | EPB | ENPOLAR | $\Delta G$ |
| --- | --- | --- | --- | --- | --- |
| Wild Type | -87.49($\pm 1.36$) | -709.22($\pm 10.19$) | 736.38($\pm 9.08$) | -10.01($\pm 0.11$) | -70.34($\pm 2.80$) |
| Omicron | -87.70($\pm 1.53$) | -1420.90($\pm 10.98$) | 1437.32($\pm 9.50$) | -9.99($\pm 0.09$) | -81.28($\pm 1.91$) |
| a475m_e484i_g476t | -94.75($\pm 1.73$) | -932.13($\pm 10.87$) | 954.70($\pm 9.26$) | -10.05($\pm 0.12$) | -82.22($\pm 3.67$) |
| e484i_g476t_l492i_p491r_t500r_y495v | -99.25($\pm 1.20$) | -1384.12($\pm 10.86$) | 1411.58($\pm 10.31$) | -10.64($\pm 0.05$) | -82.44($\pm 2.24$) |
| a475m_e484i_g476t_l492i_p491r_t500r_y495v | -103.66($\pm 1.36$) | -1402.13($\pm 12.03$) | 1434.75($\pm 11.70$) | -11.42($\pm 0.13$) | -82.46($\pm 2.82$) |
| a475m_c480h_e484i_g476t_l492i_p491r | -96.58($\pm 1.82$) | -1150.62($\pm 14.10$) | 1175.34($\pm 13.02$) | -10.82($\pm 0.11$) | -82.68($\pm 2.68$) |
| a475m_e484i_g476t_l492i_p491r_t500r | -102.97($\pm 1.53$) | -1405.63($\pm 10.42$) | 1436.23($\pm 8.83$) | -10.81($\pm 0.08$) | -83.18($\pm 2.58$) |
| a475m_c480h_e484i_g476t_l492i_p491r_t500r_y495v | -97.17($\pm 1.64$) | -1405.94($\pm 14.08$) | 1421.37($\pm 12.42$) | -11.06($\pm 0.10$) | -92.81($\pm 3.23$) |
| a475m_c480h_e484i_g476t_l492i_p491r_t500r | -102.32($\pm 1.43$) | -1418.90($\pm 11.69$) | 1438.55($\pm 10.59$) | -10.83($\pm 0.13$) | -93.51($\pm 2.86$) |

Binding free energy of multiple mutated variants. VDW: van der Waals energy, EEL: electrostatic energy, EPB: polar solvation energy, ENPOLAR: non-polar solvation energy, $\Delta G$ : binding free energy.

